# A Chemoenzymatic Method for Glycoproteomic N-glycan Type Quantitation

**DOI:** 10.1101/803494

**Authors:** Henghui Li, Leyuan Li, Kai Cheng, Zhibin Ning, Janice Mayne, Xu Zhang, Krystal Walker, Rui Chen, Susan Twine, Jianjun Li, Daniel Figeys

## Abstract

Glycosylation is one of the most important post-translational modifications in biological systems. Current glycoproteome methods mainly focus on qualitative identification of glycosylation sites or intact glycopeptides. However, the systematic quantitation of glycoproteins has remained largely unexplored. Here, we developed a chemoenzymatic method to quantitatively investigate N-glycoproteome based on the N-glycan types. Taking advantage of the specificity of different endoglycosidases and isotope dimethyl labeling, six N-glycan types of structures linked on each glycopeptide, including high-mannose/hybrid, bi-antennary and tri-antennary with/without core fucose, were quantified. As a proof of principle, the glycoproteomic N-glycan type quantitative (glyco-TQ) method was first used to determine the N-glycan type composition of immunoglobulin G1 (IgG1) Fc fragment. Then we applied the method to analyze the glycan type profile of proteins in the breast cancer cell line MCF7, and quantitatively revealed the N-glycan type micro-heterogeneity at both the glycopeptide and glycoprotein levels. The novel quantitative strategy to evaluate the relative intensity of the six states of N-glycan type glycosylation on each site provides a new avenue to investigate function of glycoproteins in broad areas, such as cancer biomarker research, pharmaceuticals characterization and anti-glycan vaccine development.

## INTRODUCTION

Glycosylation is one of the most common post-translational modifications (PTM) ^1^. Glycans exhibit vast structural microheterogeneity which is mainly generated by variable glycan structures at each of their specific glycosylation sites. The N-linked glycans are generally attached to the Asn at Asn-X-Ser/Thr consensus sequence, where X is any amino acid other than Pro^2^. The biosynthesis of N-linked glycoproteins is under a complex sequence of enzymatically catalyzed events, leading to a variety of diverse N-glycan structures. The diverse N-glycan structures are generally classified into three types: high mannose, hybrid, and complex type glycans, with all N-glycans sharing a common penta-saccharide (GlcNAc2 Man3) core structure ^3^. Although the structure of glycan is variable, evidence shows that the mammalian glycans are remarkably well conserved in certain organisms, expressing a distinct array of glycan profiles under defined conditions^4^.

Mass spectrometry (MS) is a powerful platform to comprehensively analyze protein glycosylation. However, due to the low abundance of glycosylated peptides and the heterogeneity of glycan structures, N-glycopeptide enrichment is required. Several enrichment methods have been reported, including lectin^5^ and hydrazide chemistry-based methods^6,7^, boronic acid enrichment^8^, hydrophilic interaction liquid chromatography (HILIC)^9^ and metabolic labeling^10,11^. In general, these strategies for detecting N-linked glycosylated sites require an additional de-glycosylation step by N-glycosidase F (PNGase F) before MS detection^5–7^. Unfortunately, this results in the loss of the glycan structure information at the glycosylated sites as the glycans are removed from the peptides.

Recently, a site-specific glycoproteomic method was used to detect intact glycopeptide by MS with a variety of tandem MS techniques^10,12–14^. This strategy allows the simultaneous detection of glycopeptide sequence, glycosylation site and glycan structures in one MS/MS spectrum. Current state-of-the-art MS technology with multiple dissociation known as activated ion electron transfer dissociation methods (AI-ETD) allowed intact glycoproteomic identification of more than one thousand (~1500) intact N-glycopeptides from a mouse brain tissue^13^. The site-specific glycoproteomic strategies have been applied to quantitatively detect glycoprotein alteration using the isotope labeling^12^, isobaric labeling strategies^15^ and labeling-free method^16^. However, due to missed detection of low abundant glycan structures, more than half of identified glycopeptides were linked with only one or two glycan structures using the intact glycopeptide method, hindering the comprehensive quantitation in complex biological samples^13^.

The diverse N-glycan structures play important roles in many key biological processes, including cell adhesion, receptor activation, molecular trafficking, signal transduction and disease progression, and immunotherapy^17,18^. Some apparent changes associated with cancers are the overexpression of sialylation and core fucosylation, and complex branched N-glycans. For example, increased core fucosylated type of N-glycan is an important signature of several cancers, such as hepatocellular carcinoma^19^, lung cancer^20^ and breast cancer^21^. Therefore, quantitatively monitoring the N-glycan type changes in glycosylation are important for the diagnosis, prognosis, and understanding molecular mechanisms involved in pathogenesis. Cao L et al. introduced a MS-based method that used two glycosidases PNGase F and endoglycosidase H (Endo-H), to assess the site occupancy and proportion of high-mannose and complex-type glycans of purified human immunodeficiency virus (HIV) envelope (Env) glycoprotein^22^. The NMR-based strategy was introduced to allow dissecting the glycan pattern of the IgE high-affinity receptor (FcεRIα), presenting of pauci-mannose, high-mannose, hybrid, and bi-, tri-, and tetra-antennary complex type N-glycans with different degrees of fucosylation and sialylation^23^. A purification step of glycoprotein is required in these methods, therefore, glycan type quantitation at the proteome level is urgently needed for complex biological samples.

To fulfill this analytical challenge, we developed a robust chemoenzymatic based method that quantitatively determined the proportion of N-glycan types at each glycopeptide. Briefly, three aliquots of trypsin proteolyzed sample are treated in parallel with three specific endoglycosidases and the aliquots are isotopically labeled using the three plex dimethyl labeling strategy and then combined. The cleaved N-glycopeptides are biotinylated and enriched by affinity chromatography, and the eluted N-glycopeptides are analyzed by MS. The glycoproteomic N-glycan type quantitative (Glyco-TQ) strategy was first applied on the standard glycoprotein IgG1 Fc fragment and further used to comprehensively investigate glycopeptides from the MCF7 breast cancer cell line. The data interpretation is convenient and compatible with the general proteomic platform, without need of laboriously generating sample-specific spectral libraries, complex data filtering process or specialized commercial data analysis tools. The result showed that the novel strategy could provide quantitative information on important characteristic of glycoproteins, including the relative proportion of high-mannose and linkage-related complex type glycan, and proportion of non-fucosylated and core fucosylated type glycan at each glycosylated site. To our knowledge, this is the first report to quantify the proportion of N-glycan types on the glycopeptides using the data-dependent acquisition mode for a complex biological sample.

## RESULTS

We developed a method for the quantitative analysis of N-glycan type micro-heterogeneity at both glycopeptide and glycoprotein levels. The method includes the following steps (Fig. 1a): (i) the trypsin proteolyzed peptides were divided into three aliquots, treated with one of three endoglycosidase (H, S and F3), and incubated with β-N-acetylhexosaminidasef to remove O-linked N-acetylglucosamines (O-GlcNAc); Endoglycosidase H (Endo-H) releases high mannose and hybrid type N-glycans, including those that have a fucose residue attached to the core structure^24^; endoglycosidase S (Endo-S) releases bi-antennary complex type glycans^25^; and endoglycosidase F3 (Endo-F3) release core fucosylated bi-antennary complex type glycan and tri-antennary complex glycan from N-glycopeptides^26^ (ii) the three aliquots were dimethyl labeled with ‘light’ isotope for Endo-H treated peptides, ‘intermediate’ isotope for Endo-S treated peptides and ‘heavy’ isotope for Endo-F3 treated peptides, respectively; (iii) the aliquots were combined and the retained GlcNAc on the N-glycopeptide was further transformed with N-azidoacetylgalactosamine (GalcNAz) through the catalysis of β-1,4-galactosyltransferase Y289L (GalT1 Y298L); (iv) the GalNAz labeled N-glycopeptides were covalently reacted with the photocleavable (PC) alkyne biotin through the copper(I)-catalyzed azide-alkyne cycloaddition (CuAAC) reaction; (v) the biotin linked peptides were enriched by high capacity streptavidin agarose affinity chromatography. The non-glycopeptide and reagents were removed through extensively washing; (vi) the N-glycopeptides were released by 365 nm ultraviolet irradiation and detected by MS. The detailed schematic for the PC alkyne biotin structure and reaction procedure is shown in Supplementary Fig. 1.

**Fig. 1.**
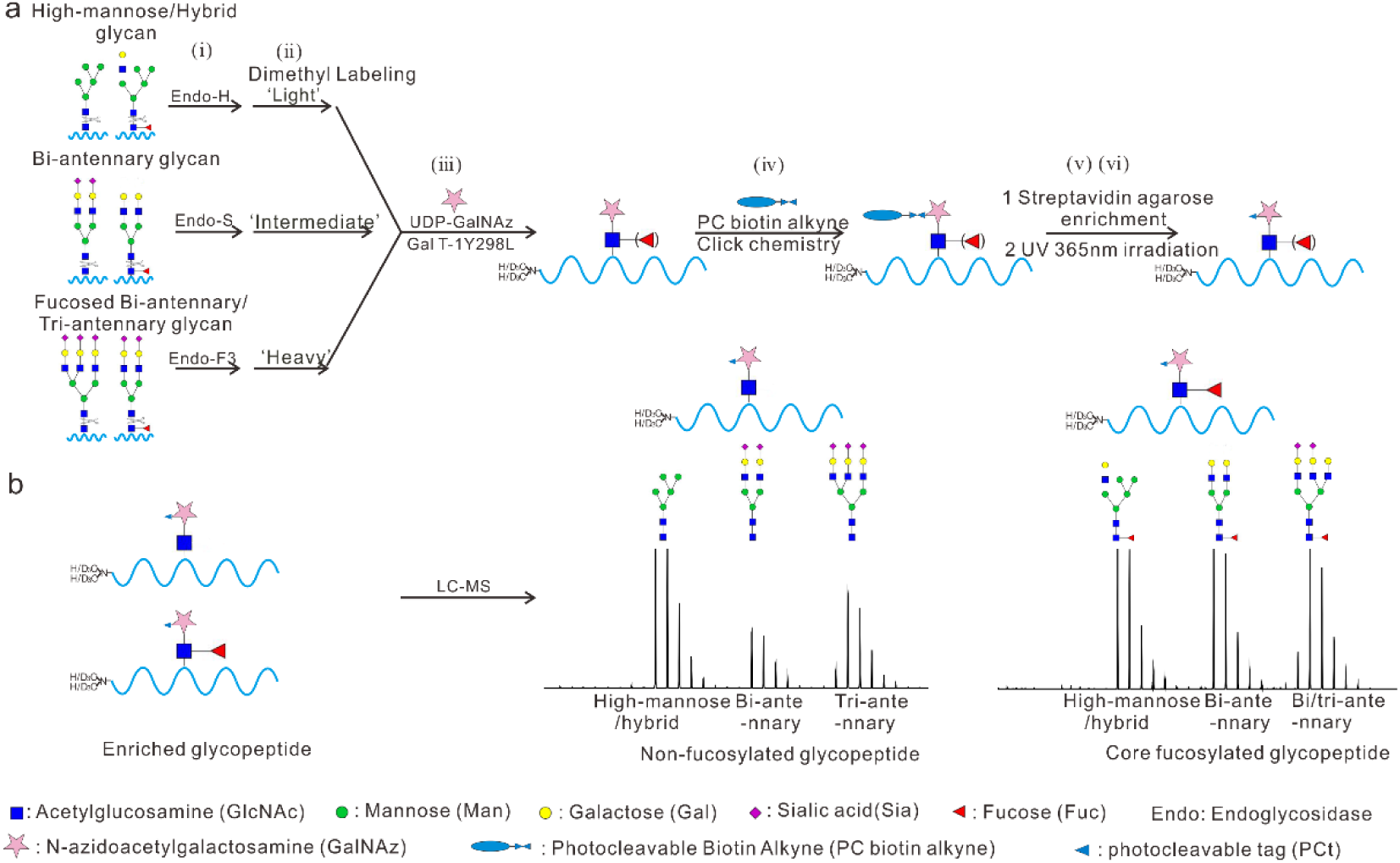
The enrichment and quantitative strategy for N-glycoproteomics. **a** The workflow for the N-glycopeptide enrichment and quantitation. **b** Six N-glycan types of glycan structures linked on the glycopeptide: non-fucosylated high mannose/hybrid type, non-fucosylated bi-antennary, non-fucosylated tri-antennary, core fucosylated high mannose/hybrid type, core fucosylated bi-antennary, core fucosylated bi/tri-antennary type.

We have tested different endoglycosidases for their specificity and selected three endoglycosidases for our method. To evaluate the specificity of the selected endoglycosidases, glycopeptides from MCF7 cells were incubated with Endo-H, Endo-S and Endo-F3, respectively. The released glycans were collected, labeled with procainamide through reductive amination and analyzed by nano LC-MS. The result showed that five high mannose and four hybrid N-glycans could be released by Endo-H (Supplementary Fig. 2a). The substrates for Endo-S were all bi-antennary complex glycans (Supplementary Fig. 2b). The Endo-F3 released the non-fucosylated tri-antennary and core fucosylated bi/tri-antennary glycans (Supplementary Fig. 2c). However, bisected bi-antennary structures with/without core fucose were not substrates for Endo-F3 (Supplementary Fig. 3). None of these three endoglycosidases showed any activity towards the more complex tetra-antennary structures. The detailed information of endoglycosidase specificity is listed in Supplementary Table 1. Therefore, using the three endoglycosidases in our method, the quantitative assessment of six N-glycan types at glycopeptides was achieved and the sex types of N-glycan were classified as following: non-fucosylated high mannose/hybrid, non-fucosylated bi-antennary, non-fucosylated tri-antennary, core fucosylated high mannose/hybrid, core fucosylated bi-antennary and core fucosylated bi/tri-antennary type glycans (Fig. 1b).

Our method also includes a novel enrichment strategy. Previously, GalT1 Y298L was used to label the O-linked β-*N*-acetylglucosamine (GlcNAc) glycopeptide with GalNAz^27^. We discovered that the GalT1 Y298L could effectively label the N-GlcNAc (on average 95.3%) as shown in Supplementary Fig. 4. After the N-linked glycopeptides were modified with GalNAz, and covalently linked with photocleavable biotin through click reaction, the unlabeled peptides and other reagents were removed by extensive washing with urea and organic buffer. The PC biotin alkyne was selected for our method development because ultraviolet irradiation is a milder condition to release the labeled glycopeptides, when compared with chemical methods that use strong reductive hydrazine or oxidizing regents^28^. Irradiation with 365 nm ultraviolet light efficiently recovered glycopeptides from the agarose streptavidin beads, with almost all glycopeptide released within 15 min (Supplementary Fig. 5).

### Validating the Glyco-TQ method on the standard glycoprotein IgG1 Fc

We first tested our approach using an immunoglobulin G1 (IgG1), containing one fragment crystallizable region (Fc fragments) and antigen-binding fragments (F(ab’)_2_ fragment). It has one fixed N-glycosylated site at Asn297 of the Fc fragment while glycan occupancy on the F(ab’)_2_ fragments are reported to be near 20%^29^. To obtain one standard N-glycoprotein with one fixed glycosylated site, IgG1 from human serum was treated with IdeS protease, to generate a homogenous pool of F(ab’)_2_ and Fc/2 fragments and then the Fc fragments were enriched by protein A agarose chromatography (Fig. 2a). As proof of principle for our quantitative method, the proportion of each glycan type was investigated by both matrix-assisted laser desorption/ionization-MS (MALDI-MS) and our novel Glyco-TQ method. The N-glycan spectrum of the Fc fragment was interrogated by MALDI-MS detection in Fig. 2b. The proportion of different types of N-glycans was calculated through the peak intensity of each glycan structure based on the MALDI-MS spectrum. The detailed ratio information of each structure is shown in Supplementary Table 2 for the MALDI-MS detection. For Glyco-TQ method, glycopeptides with sequence EEQYN#STYR were classified into six types based on their linked glycan structure: non-fucosylated high mannose/hybrid type, non-fucosylated bi-antennary type, non-fucosylated tri-antennary type, core fucosylated high mannose/hybrid type, core fucosylated bi-antennary type and core fucosylated bi/tri-antennary type glycan. The MS spectra of enriched glycopeptides and their corresponding extracted-ion chromatograms are shown in the Fig. 2c. The comparison of the proportion of glycan types based upon our Glyco-TQ method and MALDI-MS detection is shown in Fig. 2d. The high mannose and hybrid glycan type was not detected using either methods, showing the high specificity of Endo-H. As for the bi-antennary glycans released by Endo-S, the proportion of the core fucosylated bi-antennary glycan is 93.9% using our Glyco-TQ method and 90.5% using MALDI-MS detection; the proportion of the non-fucosylated bi-antennary is 6.1% using our Glyco-TQ method versus 9.5% using MALDI-MS detection. The proportion of the core fucosylated bi-antennary from our method is higher than using MALDI-MS detection (93.9% versus 90.5%), which may be due to their different ionization modes. The Endo-F3 did not release the core fucosylated bisecting bi-antennary glycans and non-fucosylated bi-antennary type glycans (showing only with minor activity when compared with tri-antennary glycans) (Supplementary Fig. 4)^30^. Only 3.3% of non-fucosylated glycans were released by the Endo-F3 in these experiments. In conclusion, our Glyco-TQ method quantitatively revealed glycan type and linkage of IgG1 Fc.

**Fig. 2.**
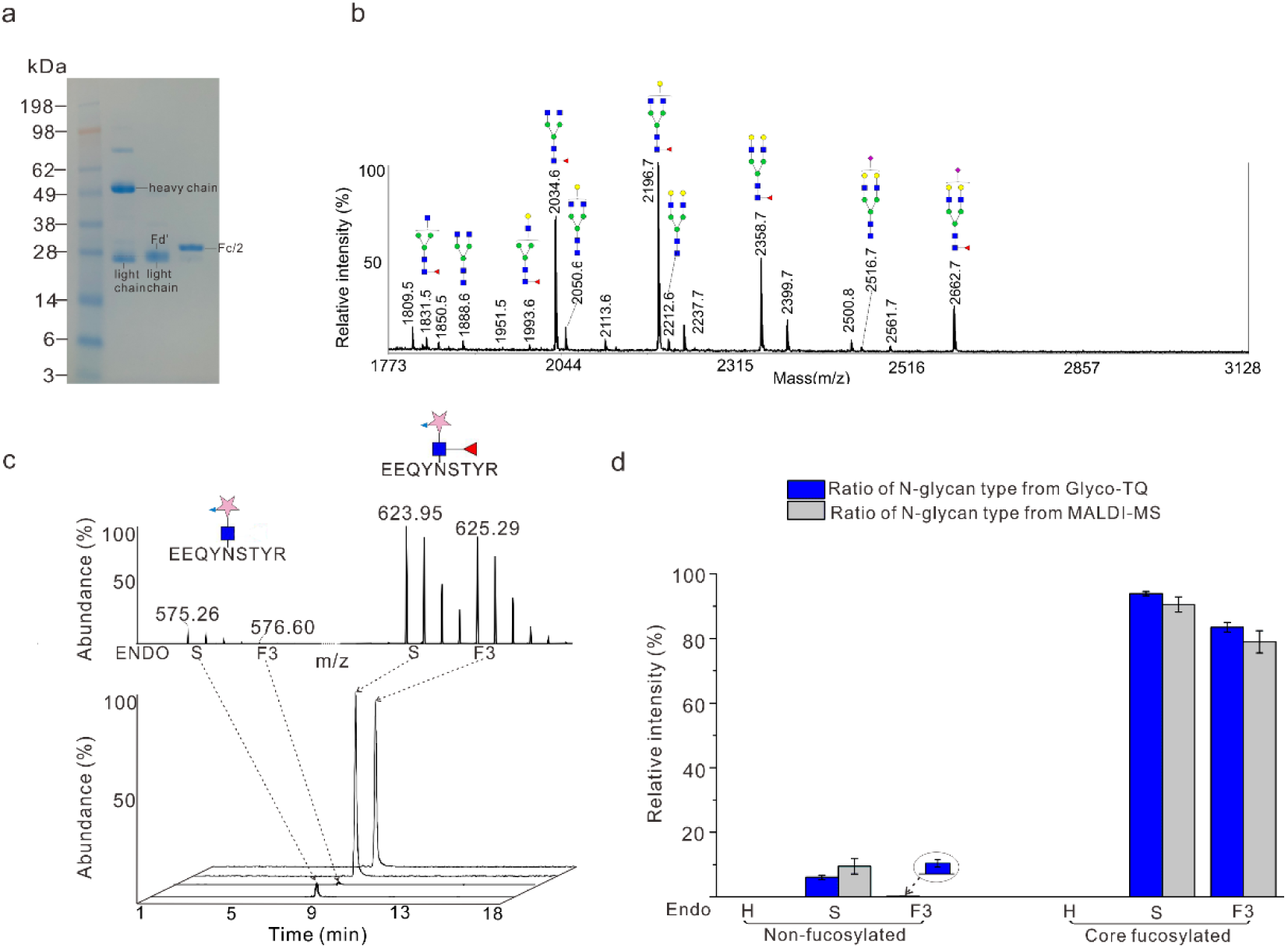
Quantitative analysis the standard glycoprotein IgG1 Fc using the Glyco-TQ method. **a** Purification of Fc fragment from human serum IgG1. **b** MALDI-MS detection the glycan profile of Fc fragments. **c** Quantitative investigation the proportion of Fc fragment glycopeptide, the MS profile of glycopeptide (up) and the corresponding extracted-ion chromatogram of glycopeptide. **d** Quantitative comparison of MALDI-MS method and Glyco-TQ method basing on the glycan type.

### Identification of N-glycopeptides from MCF7 cells

#### Detection of endogenous and native N-linked GlcNAc glycopeptides

Similar to endoglycosidases, Endo-β-N-acetylglucosaminidase (ENGase) acts as a de-glycosylation enzyme for the misfolding proteins in the cytosol^31^. We first applied our enrichment method to enrich the endogenously and native existing N-linked GlcNAc glycopeptides from MCF7 total cell lysates in absence of added any endoglycosidase (Supplementary Fig. 6). Our enrichment strategy yielded 71 N-linked glycopeptides that mapped to consensus N-glycosylated sequences (Asn-X-Ser-Thr-Cys) and that were detected with modification of one N-linked GlcNAc residue (peptide-GlcNAc) (Supplementary Table 3). At the same time, we also detected 63 O-linked GlcNAc modified peptides, which are located in the nucleus and cytoplasm by gene ontology (GO) analysis (Supplementary Table 4). Caution should be taken when interpreting results since β-N-Acetylhexosaminidasef couldn’t fully remove some abundant O-linked GlcNAc modifications.

#### Detection of N-glycopeptides from MCF7 cells

To verify our enrichment strategy, we applied the three endoglycosidases (H, S, F3) to help us enrich all the high mannose, hybrid, bi- and tri-antennary complex linked glycopeptides (Supplementary Fig. 7). To evaluate the reproducibility of our enrichment method, we performed three biological replicates with MCF-7 protein cell lysates and found 73% glycopeptides were identified in at least two replicates (Supplementary Fig. 8). We compared the glycopeptide and non-glycopeptide fractions in each parallel replicate, and showed that the specificity of our method is 55.4%. Our performance was better than the specificity of previously boronic acid and ZIC-HILIC enrichment method^8^. After N-glycan was released by the endoglycosidase, the core fucose was retained on the core GlcNAc residue, which allowed us to simultaneously distinguish the non-fucosylated and core fucosylated peptides. In total, 1090 N-glycopeptides were detected, including 916 non-fucosylated and 174 core fucosylated glycopeptide, corresponding to 504 glycoproteins (Supplementary Table 5). There were 116 glycopeptides with co-occurrence of the non-fucosylated and core fucosylated glycopeptides (Supplementary Fig. 9). As well, 58 glycopeptides were detected with only core fucosylated type glycans. Amino acid frequencies of sequences surrounding the N-glycosylated site are shown in Supplementary Fig. 10 for both the canonical and atypical N-linked glycopeptides. The above was carried out using the stepped fragmentation strategy. When the normalized collision energy (NCE) was set to 15, the fuc-GlcNAc linkage was prone to cleavage, which resulted in the parent ion and fucose neutral loss-ion as the highest peaks (Supplementary Fig. 11). For example, one MS/MS spectrum of atypical motif glycopeptide with sequence ISVN#NVLPVFDNLMQQK was identified with one core fucose as shown in the Fig. 3. In addition, the glycopeptides with common glycan tag (GlcNAc-GalNAzPCt) allowed help us to identify the glycopeptides with two glycosylated sites (Supplementary Fig. 12), overcoming significant challenges posed for their identification when using the intact glycopeptide method (5-7 KDa) ^32^.

**Fig. 3.**
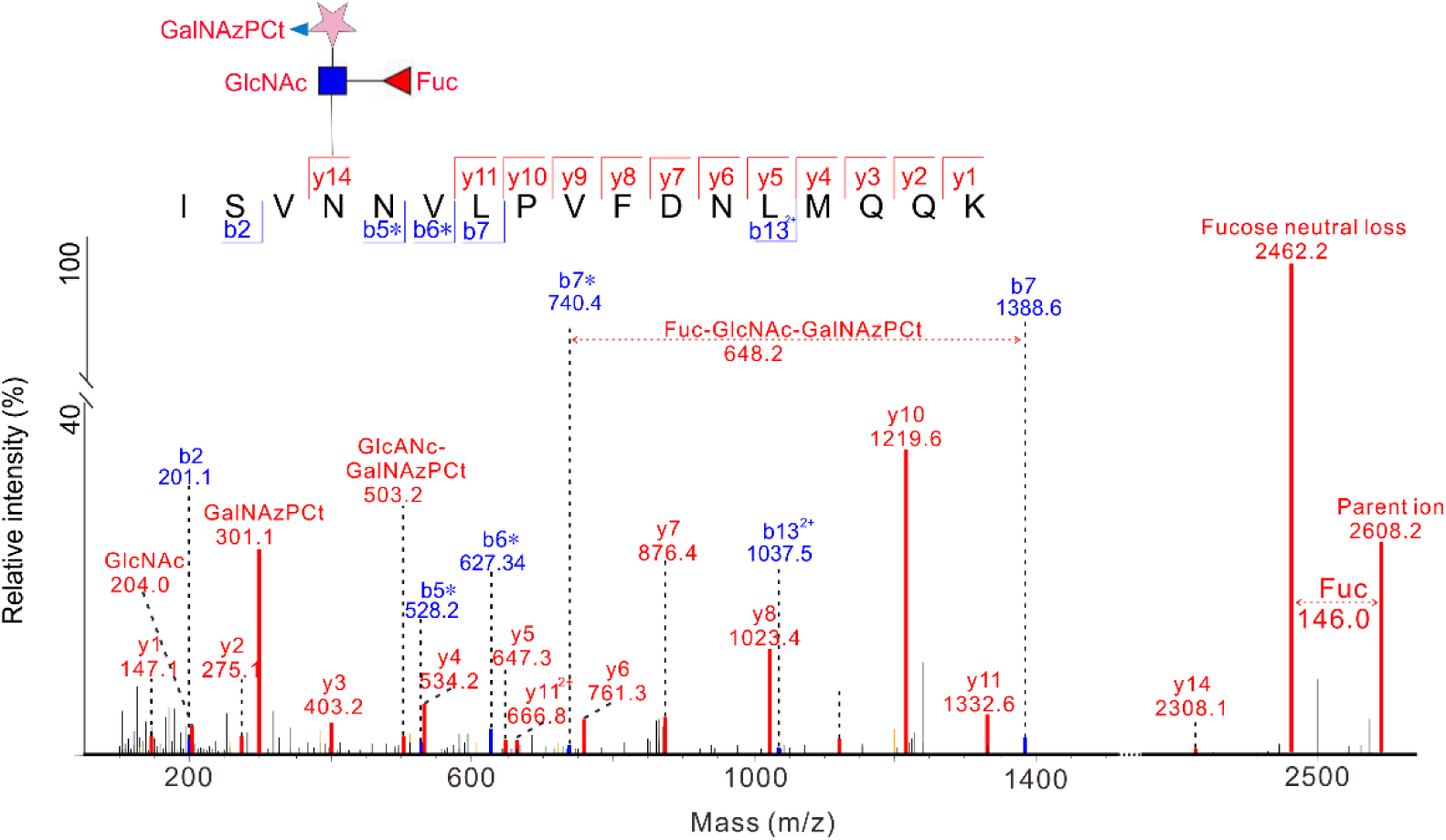
MS/MS spectrum of atypical motif glycopeptide with sequence ISVN#NVLPVFDNLMQQK. * represents the b or y ions losing the glycan common tag. The neutral loss of fucose was shown between the parent ion at m/z 2608.2 and the fucose neutral loss-ion at m/z 2462.2. The oxonium ions from the common glycan tag was set as diagnostic ion peak, representing the fragment of GlcNAz at m/z 204.0, GalNAzCA at m/z 300.1 and GlcNAc-GalNAzPCt at m/z 503.2. The mass shift between b7 and b7* (* represent losing the common tag modification) is 618.3Da, which exactly the mass of the common glycan tag (Fuc-GlcNAc-GalNAzPCt).

### Applying the Glyco-TQ method to analyze glycopeptides of MCF7 cell line

#### Quantitative micro-heterogeneity of glycopeptides and glycoproteins

The chemoenzymatic strategy allowed us to quantify the N-linked glycan dynamics on specific glycopeptide. After the peptides were treated with Endo-H, S and F3, we combined and enriched the glycopeptides by affinity chromatography (Fig. 1a). Relative quantitation of isotope peaks was calculated using the intensity of each peak area, which represented the proportion of each glycan type on the glycopeptide. First, we evaluated whether the different endoglycosidases treatment introduced biases during the glycan releasing steps. As show in Supplementary Fig. 13, high values from correlation coefficients (the value of Pearson’s correlation coefficient r > 0.96) were observed between the different endoglycosidase treatment after isotope labeling. Thus, we concluded that the endoglycosidase did not exhibit off-target protease activity and so did not introduce sample bias during glycan releasing steps. In total, we quantitatively detected 698 peptides that mapped to 378 proteins. All the glycopeptides and the relative ratio of each glycan type are shown in Supplementary Table 6. Almost all the non-fucosylated glycopeptides (> 97%) linked with the high-mannose or hybrid glycans (Fig. 4a), while only 2 glycopeptides were linked with core fucosylated hybrid glycan and more than 97% core fucosylated peptides were linked with bi- and tri-antennary complex glycans (Fig. 4b). The quantitative results exhibit distinct expression profiles for six glycan types on the glycopeptide as shown in the Fig. 4c, highlighting the abundant expression of non-fucosylated high-mannose and hybrid N-glycan. We use the heatmap to show the quantitative micro-heterogeneity of glycopeptide, and relative distribution of glycan type present on the particular glycosylated site (Fig. 4d-f). For the non-fucosylated glycopeptide, 143 glycopeptides were only linked with high-mannose and hybrid type glycan (Fig, 4d). Non-fucosylated glycopeptides (86.3%) were linked with high-mannose and hybrid type glycan, of which their proportion is larger than 50% (Supplementary Table 6 and Fig, 4d). Consistent with previous report, our results showed that N-glycans from the MCF7 cell line were predominant of non-fucosylated high-mannose/hybrid type glycans^33^. That is expected as all of the glycoproteins were first linked with high mannose type glycan during the biosynthesis^34^. As for the quantitative micro-heterogeneity on the protein level, we studied internal connection of different glycopeptides from the same glycoprotein (Fig. 4g-i). The distance between the connected dots represents the glycan type expressing divergence of glycopeptides from the same protein. Some proteins with more than one glycosylated site, such as hemicentin-1, membralin and deoxyribonuclease-2-alpha, were detected only linked with the non-fucosylated high-mannose and hybrid type glycans, released by Endo-H. However, the majority of detected proteins with multiple glycosylated sites have differential N-glycan type profiles at each site for both the non-fucosylated (Fig. 4g, h) and fucosylated glycoproteins (Fig, 4i). Due to the massive expression of non-fucosylated high-mannose and hybrid type, distribution of non-fucosylated proteins was constricted to a small region (Fig. 4g, h), while the fucosylated glycoproteins were more widely distributed, based upon the quantitative information of six N-glycan types (Fig. 4i). We also compared the location of non-fucosylated and fucosylated glycoproteins and found that more than 60% of fucosylated proteins were located in the extracellular exosome while for non-fucosylated counterparts, it was 39.8% (Supplementary Fig. 14). That result indicates the N-linked core fucose may play significant roles in cell adhesion and molecular trafficking.

**Fig. 4.**
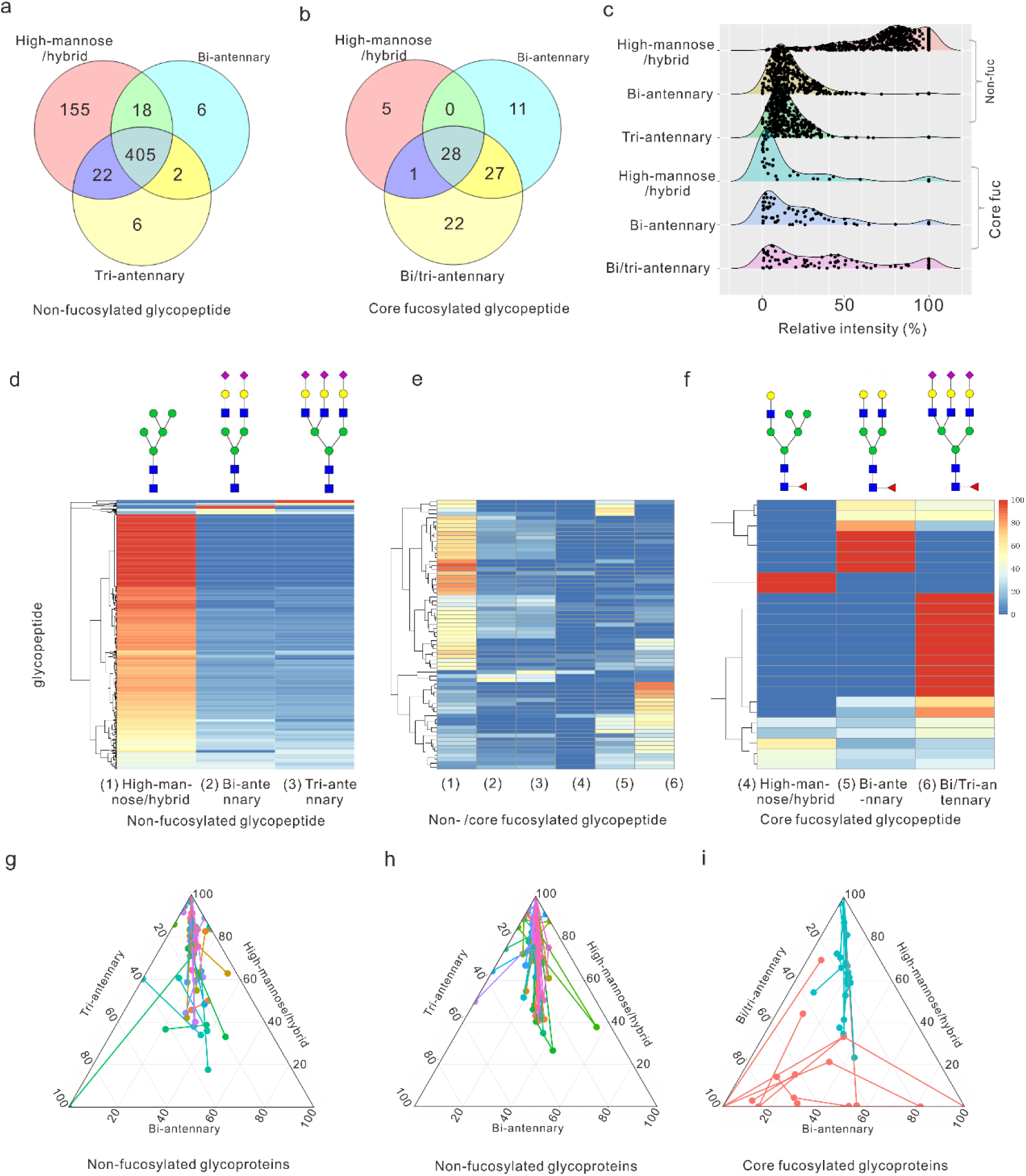
Quantitative detection MCF 7 cell derived glycopeptides by the Glyco-TQ method. **a** Detection of the non-fucosylated glycopeptide based on the glycan type. **b** Detection of the core fucosylated glycopeptide based on the glycan type. **c** The different N-glycan type ratio of glycopepide**. d** Quantitative detection of non-fucosylated glycopeptide. **e** Quantitative detection of both non-fucosylated and core fucosylated glycopeptide. **f** Quantitative detection of core fucosylated glycopeptide. Each row indicates one specific glycopeptide, and each column indicates the one type of glycan structure. The relative intensity of each glycan type on the glycopeptide was used for two-dimensional hierarchical clustering analysis. **g** The glycoprotein with two non-fucosylated glycosylation sites. **h** The glycoprotein with three or more non-fucosylated glycosylation sites. **i** The glycoprotein with two or more core fucosylated glycosylation sites, as the fucosylated glycopeptide includes non-fucosylated section and core fucosylaeted section: • represents non-fucosylated section of core fucosylated glycopeptides, • represents fucosylated section of core fucosylated glycopeptides. The distribution of each glycopeptide on the Fig. 4g-i based on the relation ratio (%) of each N-glycan type. The glycopeptides from the same glycoprotein was linked together.

#### Glycoproteins related with cancer

We quantitatively detected some glycoproteins, previously reported to be related with cancers. Mannose-6-phosphate receptor (M6PR), for example, can regulate cell growth and motility, and it functions as a breast cancer suppressor^35^. We detected seven N-glycopeptides from that protein, which exhibited diverse structures on each site (Fig. 5). The glycopeptides with sequence MN#FTGGDTCHK, TN#ITLVCKPGDLESAPVLR, N#GSSIVDLSPLIHR had similar glycan type profiles, only expressing the non-fucosylated glycan on those sites. In addition, the ratio of non-fucosylated high mannose/hybrid type glycans are more than 77% in those three peptides. The glycopeptides with sequence MDGCTLTDEQLLYSFN#LSSLSTSTFK only expressed the core fucosylated complex glycan, which could only be released Endo-F3, not Endo-S, indicating that the glycan structures were all tri-antennary core fucosylated N-glycans. The glycopeptide with sequence TGPVVEDSGSLLLEYVN#GSACTTSDGR had the most complex glycan profile, expressing both high mannose (14.9%) and fucosylated complex glycans (85.1%). The M6PR showed heterogeneity of glycan structure and distinctive glycan profiles on each of its glycopeptides. Through investigating the proportion of glycan types at each site, we could get a better understanding of the expression of this glycoprotein and its glycoprotein variants, which will promote our understanding of glycoprotein function in cancer.

**Fig. 5.**
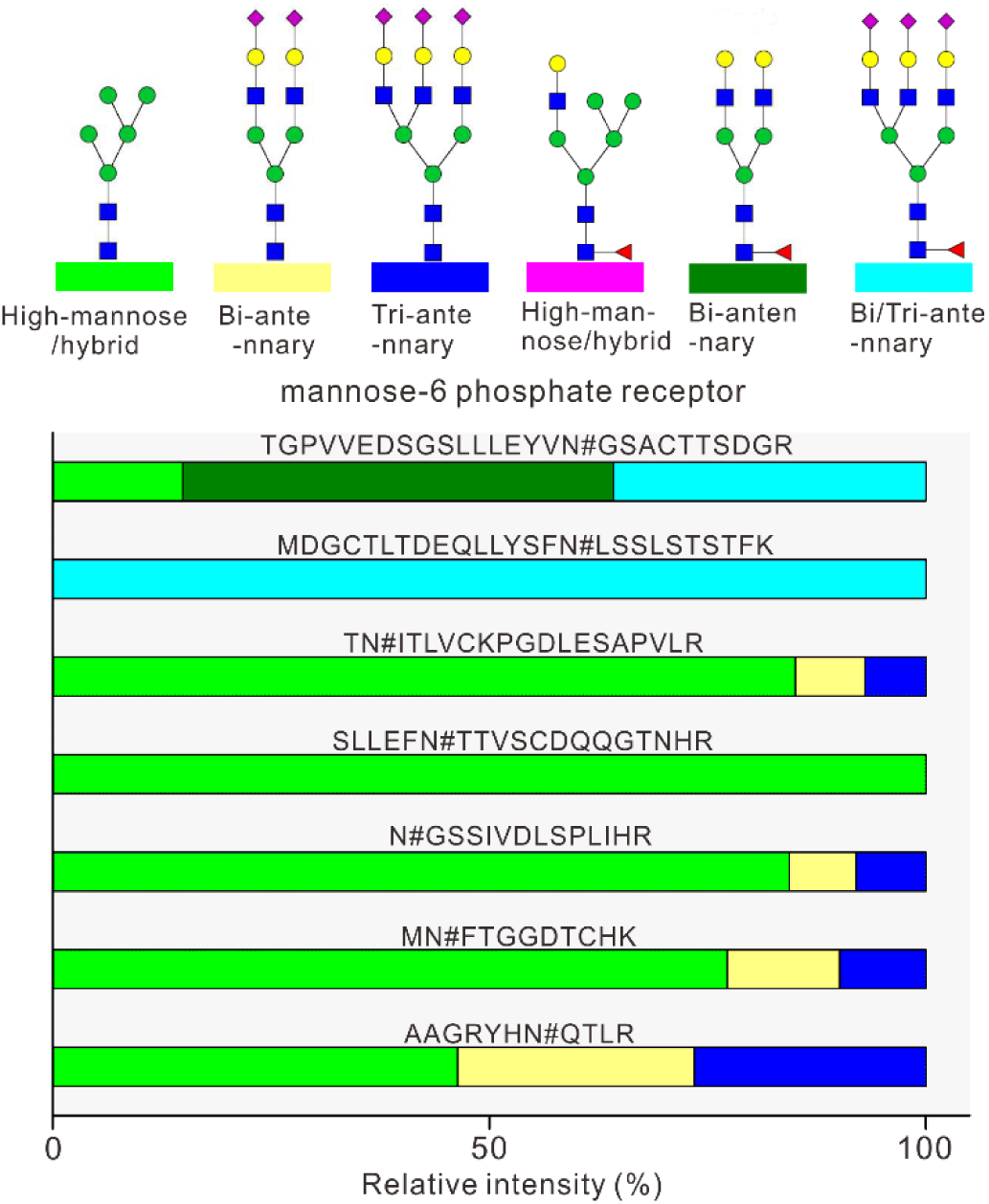
Quantitative analysis of glycosylation of mannose-6-phosphate receptor. # represents the glycosylated site.

## DISCUSSION

The previously quantitative reports of glycoproteome generally focused on the difference of one specific structure at glycopeptides between samples. The distinctive characteristic of our research is that we could provide the relative proportion of N-glycan types on each glycopeptide, which indicate the activity and aberration expression of related glycotransferases. That novel quantitative strategy provides broad information on each glycosylated site, such as the ratio of high-mannose and core fucosylated glycan, the construction of bi/tri-antennary about the complex glycans. The specific types of glycan contribute important property of glycoproteins. For example, an antibody drug linked with high-mannose type glycan showed decreased complement activity and thermal stability^36,37^. However, it is challenging to detect the low abundance N-glycans, including the high-mannose and hybrid structures^35^. We can directly provide the proportion of high-mannose and hybrid structure with help of glycol-TQ strategy, which has potential to become the routine analysis strategy for pharmaceuticals. In addition, the increasing expression of the core fucosylated type on specific glycoprotein was as potential biomarkers in some cancers. For example, core fucosylation of α-fetoprotein (AFP) L3 showed a significant increase in samples from patients with hepatocellular carcinoma (HCC) than chronic hepatitis and liver cirrhosis, and so has been approved for the early detection of HCC^19^. Taking advantage of our method, we can provide not only the expression level of fucosylated AFP L3, but also the relative ratio between non-fucosylated and core fucosylated AFP L3. Combining both level of information maybe a more sensitive and specific strategy to investigate the fucosylated biomarker. Additionally, the proteomic method could simultaneously detect multiple potential glycoproteins from single analysis. In the area of anti-glycan drug development, the N-linked glycan of HIV-1 Env is the target for broadly neutralizing antibodies, therefore, routine analysis of glycan structure supports rational design and development of vaccine immunogens^22^. Our Glyco-TQ method makes it possible to quantitatively detect glycan types on each site of human immunodeficiency virus (HIV) envelope glycoprotein (Env) trimer without extensive purification. Therefore, our Glyco-TQ method has great potential in the area of biomarker research, anti-glycan drug development and fundamental biological research.

Our strategy used relatively mild-conditions and achieved high specificity through the biotin-avidin affinity chromatography. The high specificity was contributed: the strength of the biotin-avidin binding that allows us to extensively wash to remove non-specific peptides. As well, the photocleavable tag (PCt) added to the GalNAz has one amide group, all the glycopeptides should exist in charge state of 3+ or more. The 2+ non-specific peptides wouldn’t be detected in the MS analysis, as we set the most abundant 3+ to 6+ peptides for the MS/MS analysis in the data-dependent acquisition mode. Unlike other modifications, such as phosphorylation and acetylation, the diverse glycan structures on glycoproteins make their analyses extraordinarily challenging by MS. In order to comprehensively investigate glycosylation sites, introducing a common tag could provide convenience for glycopeptide confirmation. PNGase F is the most common enzyme used to hydrolyze the glycosylamine linkage between N-glycans and asparagine, introducing a universal mass tag (0.98 Da shift) as the asparagine residue is converted to aspartic acid. However, spontaneous non-enzymatic deamidation of asparagine residues significantly affect the accuracy of the N-linked glycosylation site determination^38^. In our enrichment strategy, all processed glycopeptides contained one unique glycan residue (GlcNAc-GalNAzPCt, 502.2 Da or Fuc-GlcNAc-GalNAzPCt, 648.2 Da). That common tag not only reduced the false identification of glycopeptides, but also distinguished the non-fucosylated and core fucosylated glycopeptide. For example, the peptide labeled with Fuc-GlcNAc-GalNAzPCt, 648.2 Da was only mapped to the N-glycopeptide with a core fucose. Moreover, the common tags on the glycopeptides will help us to identify glycopeptide with atypical motifs and multiple glycosylated sites. One drawback in our research is that the HCD dissociation mode couldn’t directly tell the glycosite of glycopeptides^39^. The glycan-peptide linkage is more labile than the amino acid linkages, which leads to the core GlcNAc residue first release from peptide rather than peptide dissociation under high HCD energy. Although the N-glycopeptide canonical sequon (N-X-T/S) would help us overcome most of interference of O-linked GlcNAc modifications, that problem could be further resolved by using the more advanced EThcD dissociation strategy, which would be able to locate the modified site from the MS/MS spectrum.

Site-specific intact glycopeptide methods provide information of the exact N-glycan structure on the glycopeptide, while, the intact glycopeptides generally have lower ionization efficiency, when compared with their peptide counterparts^40^. Heterogeneity of the glycan structures from the intact glycopeptide produces a number of sub-stoichiometric modifications, splitting MS signals of the same glycopeptide into a broad spectrum of ion species^41^. Thus, the intact glycopeptide method qualitatively detects the most abundance structures for a glycopeptide, while the information of minor glycan structure on the same glycopeptide will be ignored during the MS detection. On the contrary, using our novel Glyco-TQ methods, six type structures based on the N-glycan linkage and terminal from the glycopeptide were quantified by the intensity of MS signal. Therefore, site-specific intact glycopeptide detection and our Glyco-TQ method could become complementary strategies and come together to provide both qualitative and quantitative information, facilitating further understanding of the structure and function of glycoproteins. Finally, biologists could use our strategy to directly labeling their N-glycoproteins of interest. The GalNAz labeled glycoprotein could then be modified by PEG mass tag, resolved by SDS-PAGE and visualized through immunoblotting with antibodies^42^. That strategy would permit rapid quantitation of N-glycosylation levels of particular protein without need for purification or expensive instruments, such as MS.

In conclusion, we provide a chemoenzymatic method to quantify the glycan type on the glycoproteins. All the procedures were with mild-conditions and result showed high specificity of enrichment through affinity chromatography. We provide a new quantitative strategy based the glycan type, which allows us assess the micro-heterogeneity of glycoproteins. The Glyco-TQ method has potential to be used in broad areas, such as biomarker research, pharmaceuticals, and fundamental biological research.

## METHODS

### Materials

Endoglycosidase S, F3, N-glycosidase F (PNGase F), β-N-Acetylhexosaminidasef and alkaline phosphatase were from New England Biolabs. Mutant β1-4-Galactosyltransferase (Gal-T1 Y289L)) and High Capacity Streptavidin Agarose were obtained from Thermo Scientific. IdeS protease was purchased from Promega. UDP-GalNAz was from chemily Glycoscience. The immunoglobulin G1 (IgG1) from normal human plasma was obtained from Athens Research & Technology. 2-(4-((bis((1-(tert-butyl)-1H-1,2,3-triazol-4-yl) methyl) amino) methyl)-1H-1,2,3-triazol-1-yl) acetic acid (BTTAA), photocleavable biotin alkyne (PC biotin alkyne) were purchased from Click Chemistry Tools. EDTA-free protease inhibitor cocktail was obtained from Roche Diagnostics. RapiGest SF surfactant and the Sep-Pak tC18 cartridge was obtained from Waters. All other chemical materials, if not special highlighted, was obtained from Millipore Sigma.

### Preparing the standard glycoprotein

Immunoglobulin G1 (100 μg) was treated with 1000 units IdeS protease for 3 hours in 1X phosphate buffered solution (PBS, pH 7.6). 200 µL of immobilized Protein A resin slurry (50% w/v) was added to the reaction buffer, and incubated with gentle mixing for 2 hours at room temperature. Then the Protein A resin slurry were transferred into centrifuge columns and Protein A resin was washed with 1XPBS three times to remove unbound F(ab’)_2_ fragments (fragment antigen-binding). The Fc fragments (fragment crystallizable region) of immunoglobulin G1 (IgG 1) were eluted with 100 mmol/L glycine buffer, pH 2-3. The Fc fragments were immediately neutralized with 1 M Tris-HCl buffer and stored in −80 °C for further use.

### Cell culture, protein extraction and protein digestion

The MCF-7 cell line was obtained from American Type Culture Collection (ATCC). MCF-7 cells were maintained in advanced MEM media (Gibco) with 10% (v/v) FBS, 1X GlutaMAX (Gibco), and 2.8 µg/mL Gentamicin (Gibco). The cells were cultured at 37 °C and 5% CO2. Once the cells reached 80% confluency, cells were harvested in the ice-cold RIPA buffer (50 mM HEPES pH 7.6, 150mM NaCl, 1% NP-40, 1% sodium deoxycholate, 0.1% sodium dodecyl sulfate (SDS) and protease inhibitor cocktail (CompleteMini, Roche)) by scraping. Cell lysates were subjected to ultrasonicate (10 s process with 10 s interval for 1 min) on ice using a Q125 Sonicator with 50% amplitude. The cell debris was removed through centrifugation at 16000g, 4 °C, 10 min. The protein in the supernatant was precipitated using 6-fold volume ice cold acetone overnight at −20 °C. Protein was pelleted by centrifugation at 16000g, 4 °C, 10 min and washed with the ice-cold acetone two times. For the in-solution trypsin digestion, the procedure was performed as the previously report^43^. Briefly, the resulting protein was dissolved in 50 mM ammonium bicarbonate and 8 M urea solution, reduced in 5 mM dithiothreitol (DTT) (56 °C, 30 min), and alkylated by with 10 mM iodoacetamide (25 °C, 40 min in the dark). Cell proteins were digested with the protein: trypsin (Worthington Biochemical Corp) at ratio, 50:1, in 50 mM ammonium bicarbonate, 1M urea solution pH 7.8 at 37 °C for 20 hours. After the digest, the solution was acidified (pH 2-3) by 0.5% formic acid and centrifuged to remove the debris, the supernatant was collected and peptide was desalted by Sep-Pak tC18 cartridge (Waters). The peptide elution was dried by SpeedVac concentrator (Thermo Scientific).

### Endoglycosidases digestion and dimethyl labeling

To leave one N-acetylglucosamine for high mannose linked N-peptide, peptide from 1 mg protein was parallelly digested with 0.05 U Endo-H (sigma) in the 50 mM sodium acetate buffer, pH 6; 2000 units Endo-S (NEB) for biantennary N-glycan in 50 mM sodium acetate, 5 mM calcium chloride buffer pH 5.5; 100 units Endo-F3 (NEB) in 50 mM sodium acetate, pH 4.5, for 24 hours respectively. After the endoglycosidase were denatured at 95 °C, 5 min, β-N-acetylhexosaminidasef was added to remove O-GlcNAc for another 8 hours. After desalted by Sep-Pak tC18 cartridge, the peptide was adjusted to pH 6 with HEPES buffer. For isotope dimethyl labeling, the peptides were treated with 10 μL 20% (v/v) CH2O, 15μL 3M NaBH3CN for the Endo-H treated sample; 10 μL 20% (v/v) CD2O, 10 μL 3M NaBH3CN for the Endo-S treated sample; 15 μL 20% (v/v) CD2O, 15μL 3M NaBD3CN for the Endo-F3 treated sample, at 25 °C for 45 min with mixing. The reaction was quenched by adding 10 μL 20% (v/v) ammonia solution, combined, purified by Sep-Pak cartridge, and dried by Speedvac.

### Glycopeptide enrichment

All the peptide was resuspended in the HEPES buffer (pH 7.9) containing 5 mM Zn^2+^, 2 μL phosphatase, 10 μL GalT1 T298L/1 mg peptide, and 25 μg UDP-GalNAz/1 mg peptide, incubated in 4 °C for 24 hours. Excess UDP-GalNAz was remove by Sap-pak C18 cartridge. The peptide was dried in the speedVac and resuspended the PBS buffer. The Copper(I)-catalyzed Azide-Alkyne Cycloaddition (CuAAC) reaction reagents (25 nmol PC biotin alkyne, 300 μM CuSO4, 600 μM BTTAA, 1.50 mM sodium ascorbate) was mixed with the GalNAz labeled peptide and the reaction was incubated for 3 h at 25 °C. 200 μL high capacity streptavidin agarose resin was add to the mixture and incubate overnight at 4 °C. The beads were extensively washed with 2M urea ten times, 1XPBS buffer (pH 7.6) ten times and 20% (v/v) acetonitrile (ACN) ten times. The beads were then resuspended in 50% (v/v) ACN, transferred to clear thin-walled polymerase chain reaction (PCR) tubes, and illuminated by 365 nm UV (VWR transilluminator, LM-20E) for 30 min at 4 °C with gentle mixing. The supernatant from each fraction was collected, lyophilized, and stored at −20°C.

### Mass spectrometry analysis

Glycopeptides were analyzed by an Eksigent nanoLC liquid chromatograph that was connected in-line with an Q Exactive HF-X MS. The separation of peptides was performed on an analytical column (75 μm × 50 cm) packed with reverse phase beads (1.9 μm; 120-Å pore size; Dr. Maisch GmbH) with 2-hour gradient from 5 to 35% acetonitrile (v/v) at a flow rate of 200 nl/min. The full scan mass spectrums were acquired over range 300-1800 (m/z) with the mass resolution setting 70000 at m/z 400. Maximum injection time 100 ms; AGC target value 1e6. The 12 most intense ions were selected for tandem mass spectrometry detection with the following parameters: collision energy, 30%; exclusion ions charge 1, 2, 7, 8, >8; resolution 17500, AGC target 1e5; maximum injection time 120 ms.

### Data analysis

The raw data were processed using the MaxQuant software and searched against with UniProt human database containing all proteins in the UniProt Human (Homo sapiens) database (20190802). The general parameters were performed during the search: 10 ppm precursor mass tolerances; digested with trypsin; two max missed cleavages; fixed modifications: carbamidomethylation of cysteine (+57.0214); variable modifications: oxidation of methionine (+15.9949). The common tag was also performed as variable modifications: modified amino acid, asparagine (N); composition H30C19N6O10, GlcNAc-GalNAz-photocleavable tag (GlcNAc-GalNAzPCt), 502.2023 Da; neutral losses, GlcNAc-GalNAzPCt, GalNAzPCt H17C11N5O5; diagnostic peaks, GalNAzPCt H17C11N5O5, GlcNAc H13C8NO5, GlcNAc-GalNAzPCt. If the N-glycopeptide was modified with core fucose, fucosylated linked was performed as variable modifications as following: modified amino acid, asparagine (N); composition H40C25N6O14, Fuc-GlcNAc-GalNAzPCt, 648.2602 Da; neutral losses, Fuc H10C6O4, Fuc-GlcNAc-GalNAzPCt, GalNAzPCt, Fuc+GalNAzPCt H27C17N5O9; diagnostic peaks, GalNAzPCt, GlcNAc, Fuc, GlcNAc-GalNAzPCt, Fuc-GlcNAc-GalNAzPCt and Fuc-GlaNAzPCt H23C14NO9.

## Acknowledgements

This work was supported by the Government of Canada through Genome Canada and the Ontario Genomics Institute (OGI-114), CIHR grant (ECD-144627), the Natural Sciences and Engineering Research Council of Canada (NSERC, grant no. 210034), the Ontario Ministry of Economic Development and Innovation (REG1-4450) and The University of Ottawa. DF acknowledges a Distinguished Research Chair from the University of Ottawa.

## Authors’ contributions

D.F., J.L, and H.L. designed the study. H.L., L.L, K.C and K.W performed the experiments and data analysis. R.C and S.T involved in discussion of the study design. D.F., J.L, H.L., X.Z, Z.N and J.M. wrote the manuscript. All authors participated in the data interpretation, discussion and edits of the manuscript.

