## Supplementary Note and Figures for "A Chemoenzymatic Method for Glycoproteomic N-glycan Type Quantitation"

### METHOD

#### N-glycan detection

The standard glycoprotein IgG1 Fc fragment (50 μg) was dissolved in 50 μL 50 mM HEPES buffer (pH8.0) containing 1% RapiGest SF surfactant, 50 mM dithiothreitol (DTT), 0.1% (w/v) SDS and denatured at 95 ºC for 5 min. The protein was subsequently cooled down to 50 °C and 1 μL of Rapid PNGase F was added to the protein solution. The releasing reaction was performed at 50 °C with gently mixing, Sequentially, TMPP-Ac labeling and purification were performed as previously report[1]. Briefly, a freshly prepared solution of 1mg TMPP-Ac-OSu in acetonitrile (50 μL) was added to the reaction tube with agitated vigorously agitation for 30 min at room temperature. Purification was performed by an HILIC method using home-packed microcrystalline cellulose SPE as follows: the HILIC cartridge was first washed with 3.0 mL of water and then equilibrated with 3.0 mL binding solution of ACN/H2O/acetic acid (80:17:3, v/v/v). The reaction solution was diluted in 500 μL binding solution and added to the cartridge. Finally, the cartridge was washed with 3.0 mL binding solution to remove the impurities. Derivatized glycan was eluted by 1 mL of ethanol/H2O (1:1, v/v) and dried by a Speedvac concentrator. All of the MALDI-TOF-MS detection were performed using 4800 MALDI-MS (AB SCIEX) equipped with a 355 nm Nd: YAG laser in the reflector positive mode. CHCA matrix was prepared in 50% ACN aqueous solution with a final concentration of 5 mg/mL. Samples of 0.5 μL mixed with 0.5 μL freshly prepared CHCA matrix were directly loaded onto the stainless steel MALDI plate and allowed to be dried at room temperature.  A total of 1000 laser shots were employed in each sample spot. The data processing was further analyzed by Data Explorer 4.0 (AB SCIEX).

For analysis the glycan from MCF 7, peptide from 1 mg protein was treated with Endo-H，Endo-S，Endo-F3 and PNGsee F, respectively. All peptides were boiled for 5 min to de-activate the activity of enzyme, and the released glycans were purified using PGC cartridges as previously report. PGC SPE cartridges were washed with 3.0 mL 80 % (v/v) ACN containing 0.1 % TFA and equilibrated with 3.0 mL 5 % (v/v) CAN, 0.1 % TFA. The released glycans were resolved in 5 % (v/v) ACN containing 0.1 % TFA and loaded onto the PGC cartridges. The PGC cartridge was washed with 3 mL 5 % (v/v) ACN, 0.1 % (v/v) TFA. Finally, glycans were eluted with 40 % (v/v) ACN in 0.1 % (v/v) TFA (1.0 mL). The fraction was collected and dried by Speedvac concentrator. The collected glycans were mixed with 50 μL freshly prepared labeling solution (271.7 mg/mL procainamide hydrochloride and 10.7 mg 2-picoline-borane in DMSO containing 30% (v/v) glacial acetic acid)[2]. The mixture was incubated at 65°C for 2 h.

Purification was performed by an HILIC method using home-packed microcrystalline cellulose SPE as above description. The procainamide labeled N-glycan was analyzed by an Eksigent nanoLC liquid chromatograph that was connected in-line with an Q Exactive mass spectrometer. The separation of N-glycan was performed on an analytical column (75 µm × 50 cm) packed with reverse phase beads (1.9 µm; 120-Å pore size; Dr. Maisch GmbH) with 40 min from 5 to 35% acetonitrile (vol/vol) at a flow rate of 200 nl/min. The full scan mass spectrums were acquired over range 300-1800 (m/z) with the mass resolution setting 70000 at m/z 400. Maximum injection time 100 ms; AGC target value 1e6. The 12 most intense ions were selected for tandem mass spectrometry detection with the following parameters: collision energy, 20%; exclusion ions charge 1, 5, 6, 7, 8, >8; resolution 17500, AGC target 1e5; maximum injection time 120 ms.

**SFig. 1.** The detail information of the Glyco-TQ method.

**SFig. 2.** Detection the specificity of endoglycosidase.

**SFig. 3.** The glycan structures could not be released by the endoglycosidase F3.

**SFig. 4.** The ion-extracted chromatogram of glycopeptide after labeled with GalNAz.

**SFig. 5.** The glycopeptide releasing efficiency form streptavidin agarose with 365 nm ultraviolet irradiation.

**SFig. 6.** The enrichment workflow for the endogenous and native N-linked GlcNAc glycopeptide from MCF 7.

**SFig. 7.** The enrichment workflow for the N-linked glycopeptide from MCF 7.

**SFig. 8.** Comparison of protein N-glycosylation sites identified in MCF7 cells in three biological replicate experiments.

**SFig. 9.** Overlapping of non-fucosylated and core fucosylated glycopeptide.

**SFig. 10.** Distribution and preference of glycosylated consensus sequence derived using Weblogo.

**SFig. 11.** MS/MS spectra of EEQYN#STYR with different collision energies.

**SFig. 12.** MS/MS spectrum of glycopeptide (VIN#ETWAWKN#ATLAEQAK) with two glycosylated sites.

**SFig. 13.** Evaluation the influences of different endoglycosidase on the sample.

**SFig. 14.** The comparison cellular compartment distributions of non-fucosyled glycoproteins and core fucosylated glycoproteins.

**STable 1.** The N-glycan detected from MCF 7 and substrates for the endoglycosidases (H, S, F3).

**STable 2.** The N-glycan detected from IgG1 Fc fragments, and substrates for the endoglycosidase (H, S, F3).

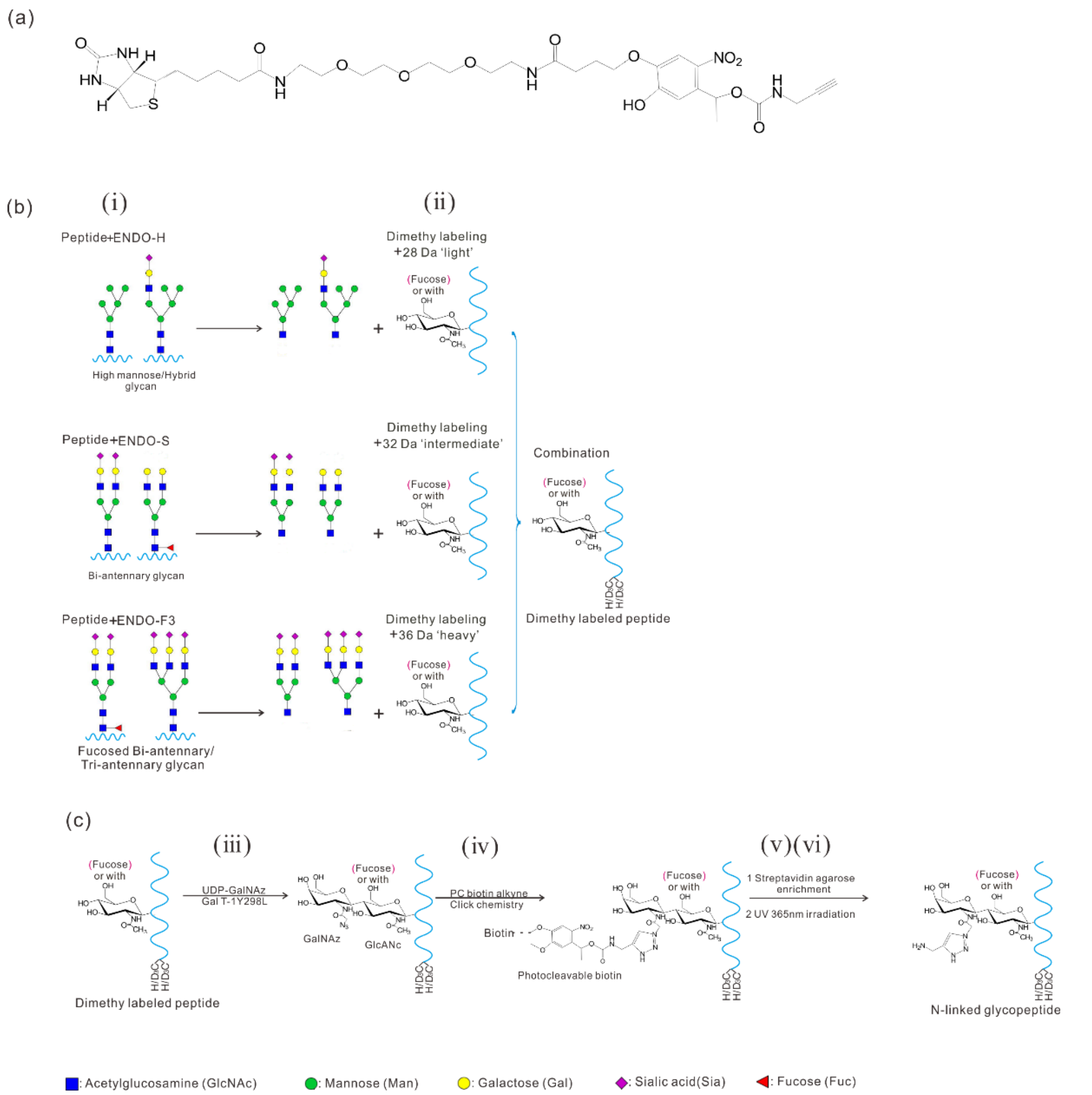


**SFig.1.** The detail information of the Glyco-TQ method. (a) the structure of photocleavable (PC) biotin alkyne; (b) procedure of endoglycosidase releasing and dimethyl labeling; (c) the process of GalNAz labeling, click reaction with PC alkyne and 365 nm ultraviolet (UV) light releasing.

**SFigure 1**

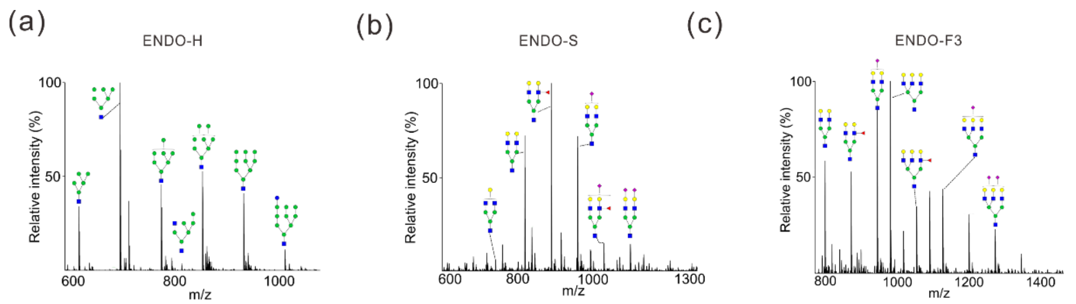


**SFig. 2.** Detection the specificity of endoglycosidase. (**a)** The glycan structure released by Endo-H. (**b)** The glycan structure released by Endo-S. (**c)** The glycan structure released by Endo-F3.

**SFigure 2**

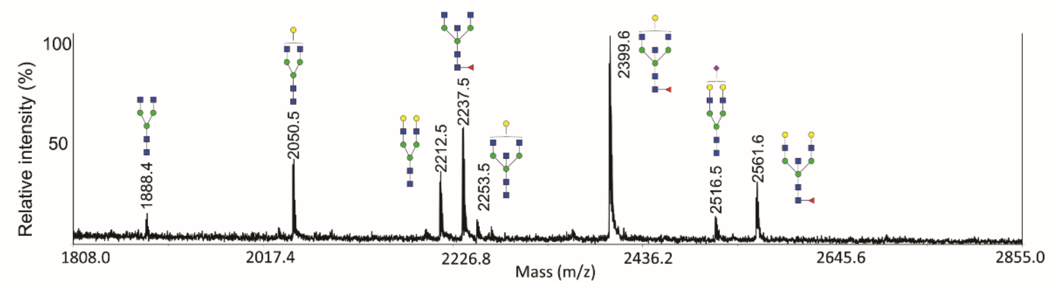


**SFig. 3.** The glycan structure could not be released by the endoglycosidase F3. After the N-glycan was rapidly released by PNGase F and labeled with TMPP-Ac-OSu. Then TMPP labeled N-glycan was treated with Endo-F3 overnight, enriched by HILIC SPE, and detected by MALDI-MS. The result showed that the Endo-F3 could not releasing the non-fucosylated biantennary and bisecting type N-glycan.

**SFigure 3**

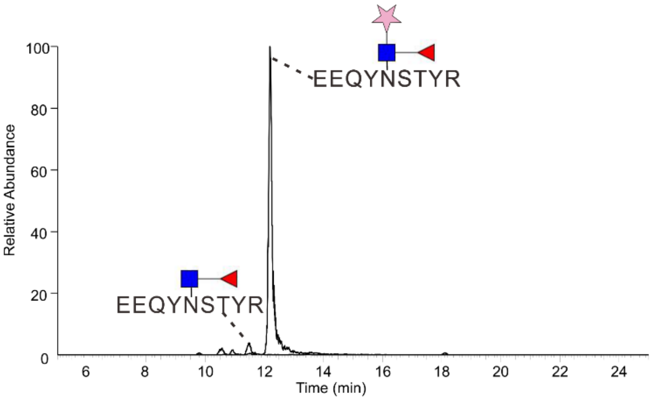


**SFig. 4.** The ion-extracted chromatogram of glycopeptide after labeled with GalNAz.

**SFig. 4**

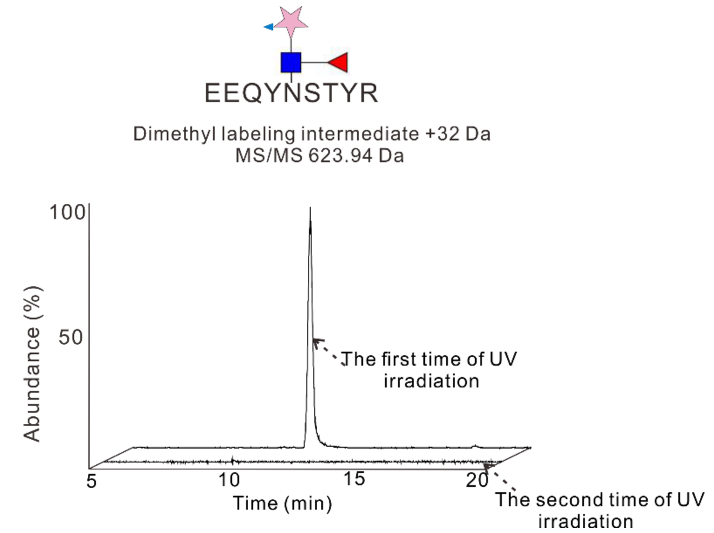


**SFig. 5.** The glycopeptide releasing efficiency form streptavidin agarose with 365 nm ultraviolet irradiation. The streptavidin agarose linked with the glycopeptides was irradiated with 365 nm UV light for 15 min, then the glycopeptides were collected. The streptavidin agarose was washed two times with water and irradiated with 365 nm UV light, 15 min for the second time. The result showed that almost all the glycopeptides was released with the first 15min UV irradiation.

**SFigure 5**

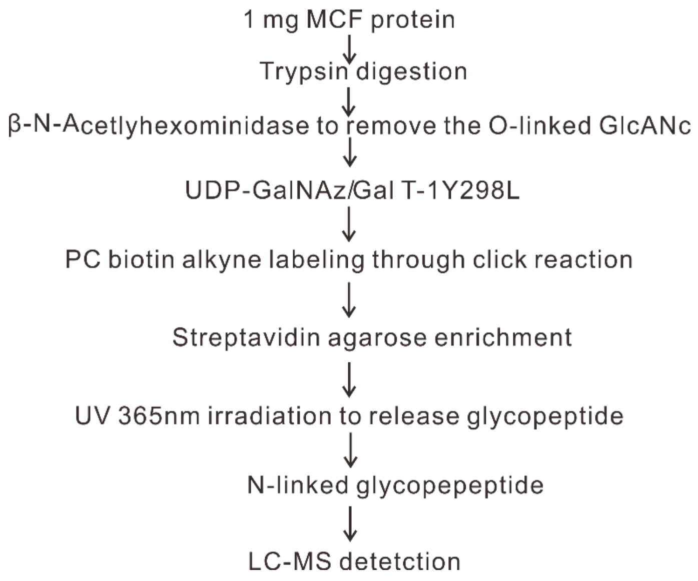


**SFig. 6.** The enrichment workflow for the endogenous and native N-linked GlcNAc glycopeptide from MCF 7.

**SFigure 6**

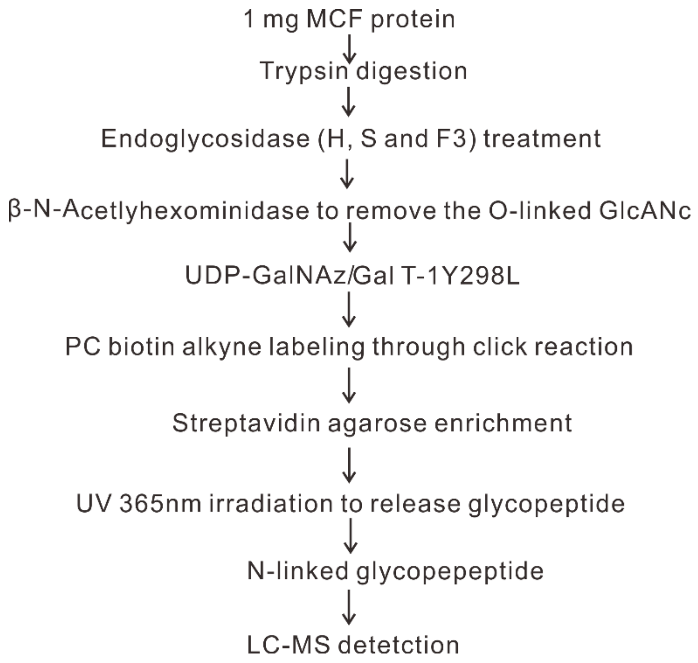


**SFig. 7.** The enrichment workflow for the N-linked glycopeptide from MCF 7.

**SFigure 7**

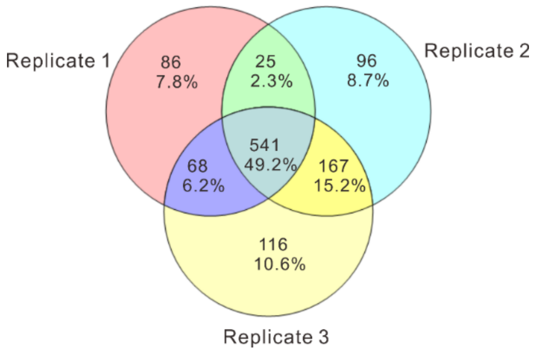


**SFig. 8.** Comparison of protein N-glycosylation sites identified in MCF7 cells in three biological replicate experiments

**SFigure 8**

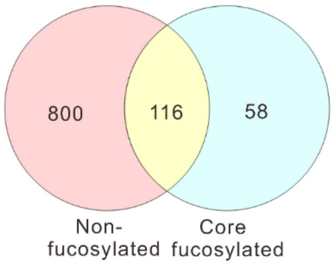


**SFig. 9.** Overlapping of non-fucosylated and core fucosylated glycopeptide.

**SFigure 9**

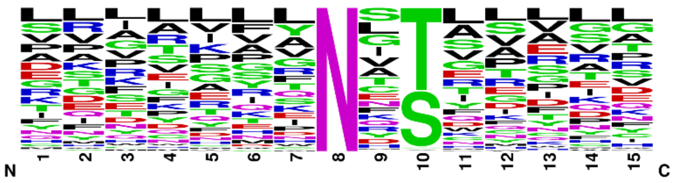


**SFig. 10.** Distribution and preference of glycosylated consensus sequence derived using Weblogo.

**SFigure 10**

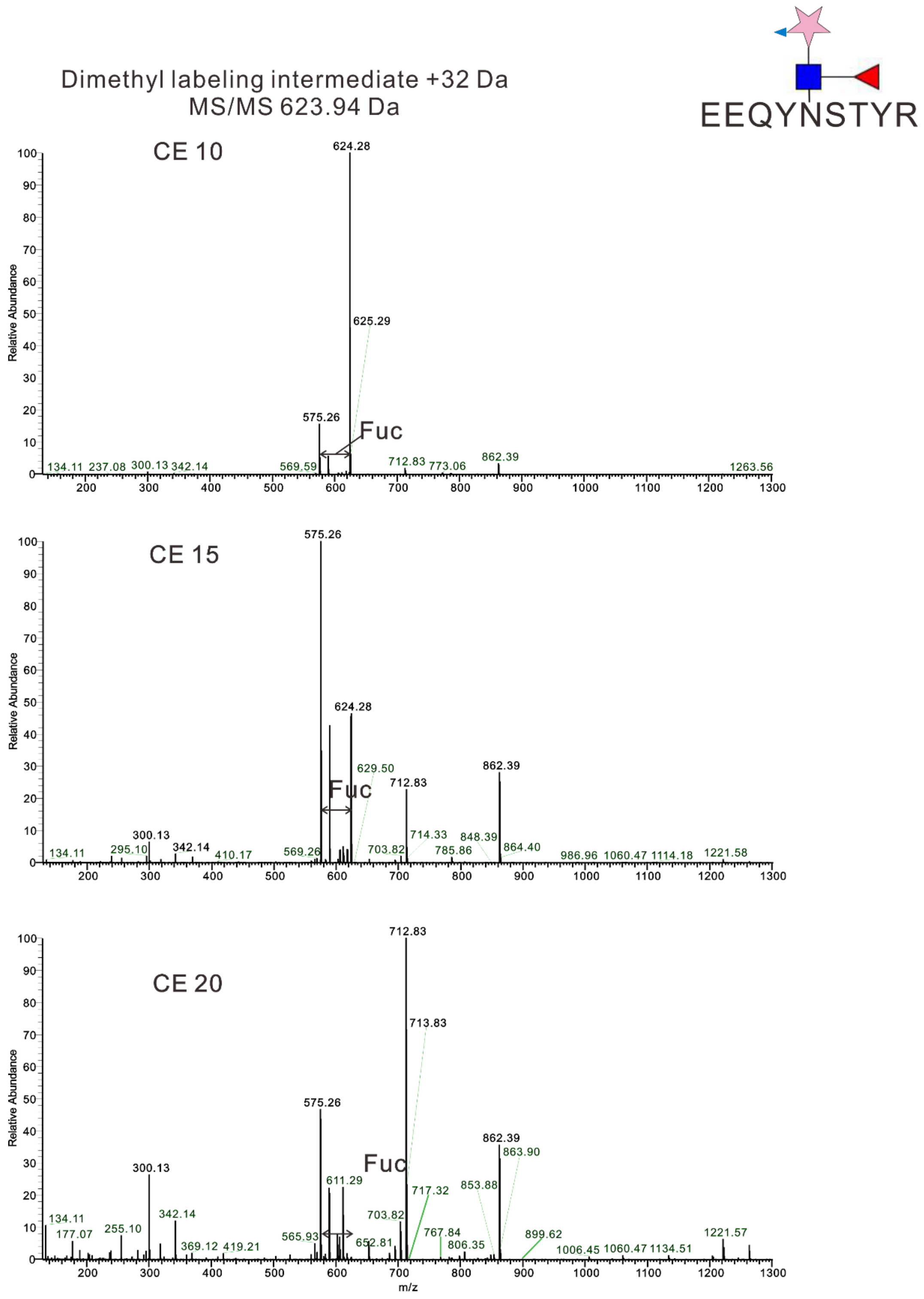


**SFig. 11.** MS/MS spectra of EEQYN#STYR with different collision energies: (a) NCE = 10; (b) NCE = 15; (c) NCE = 20. When the NCE was at 15, both the parent ions and fucose-loss neutral ions were the highest peaks in the ms/ms spectrum. The mass shift of fucose help determine the fucosylated glycopeptide.

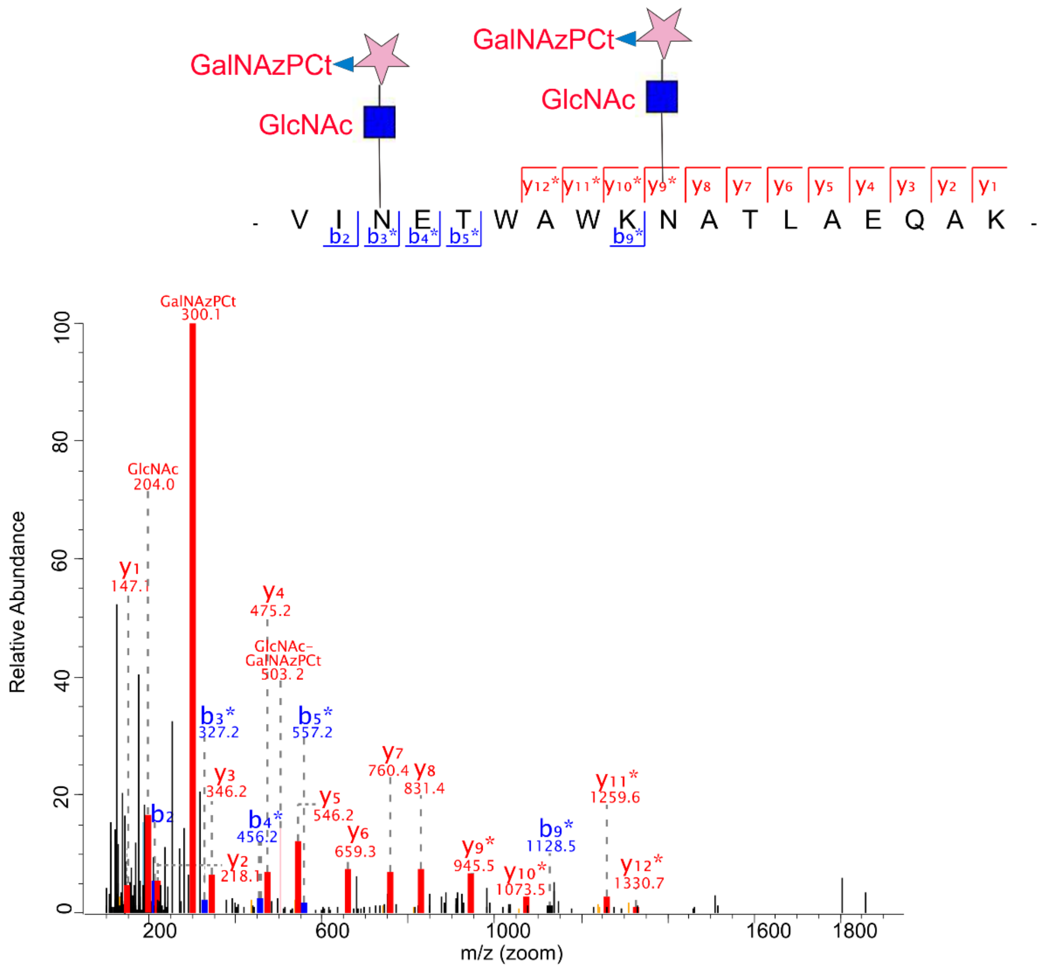


**SFig. 12.** MS/MS spectrum of glycopeptide (VIN#ETWAWKN#ATLAEQAK) with two glycosylated sites, * represents the b or y ions losing the glycan common tag. the glycopeptide with sequence VIN#ETWAWKN#ATLAEQAK has two canonical sequon (NET and NAT). The parent ion of the glycopeptide is 3076.4 Da, and the peptide backbone is 2072.0 Da. The mass difference between parent ions and peptide backbone is 1004.4 Da, which is exactly the mass of two GlcANc-GalANzPCt modification. Therefore, the glycopeptides with two special glycan tag (GlcANc-GalANzPCt) will help us identify the glycopeptide with more than one glycosylated site.

SFigure 12

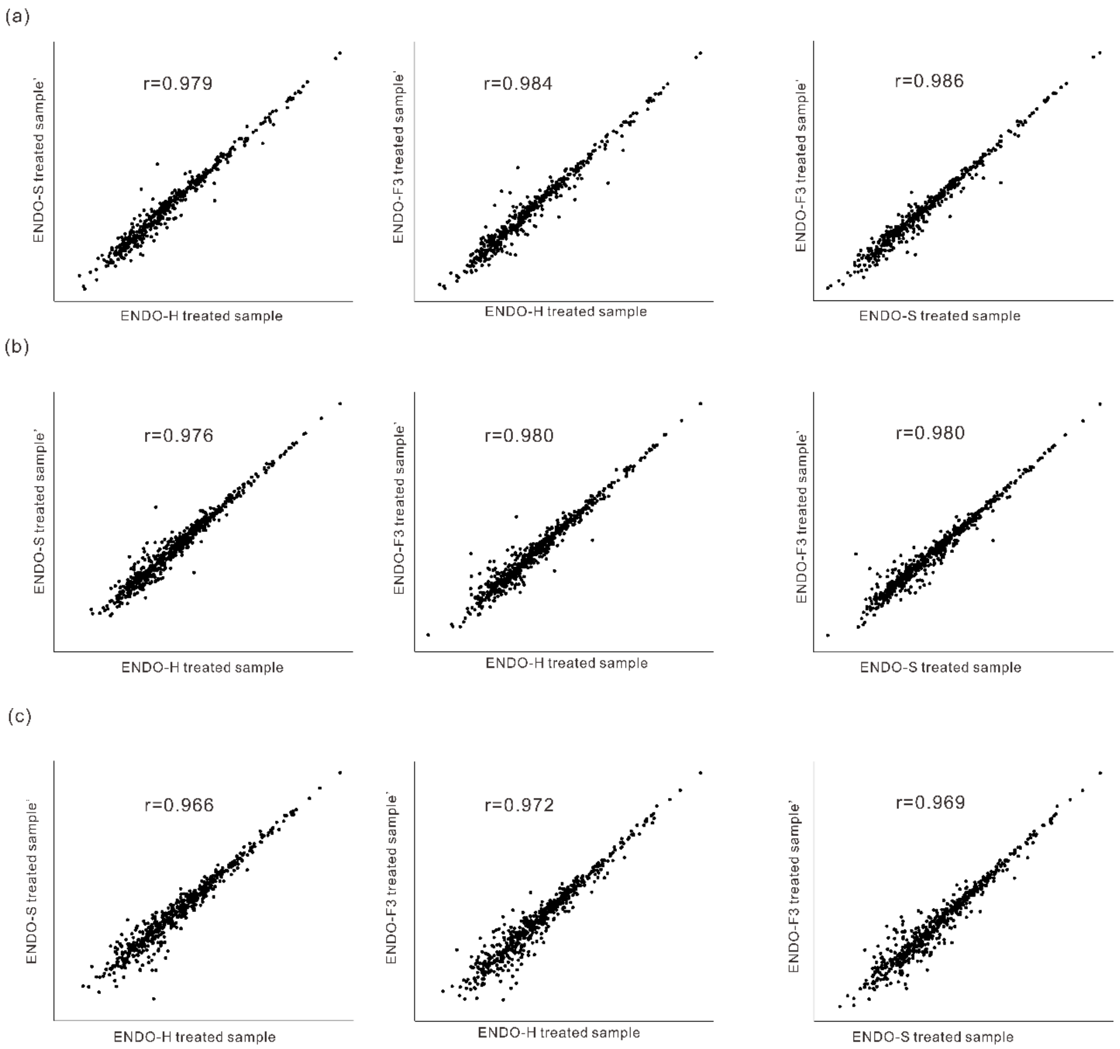


**SFig. 13.** Evaluation the influence of different endoglycosidase on the samples. The different isotope labeled sample after endoglycosidase treatment showed high degrees of correlation. Scatter plots were plotted with Log10 transformed LFQ protein intensity. Pearson's correlation coefficient r is shown.

**SFigure 13**

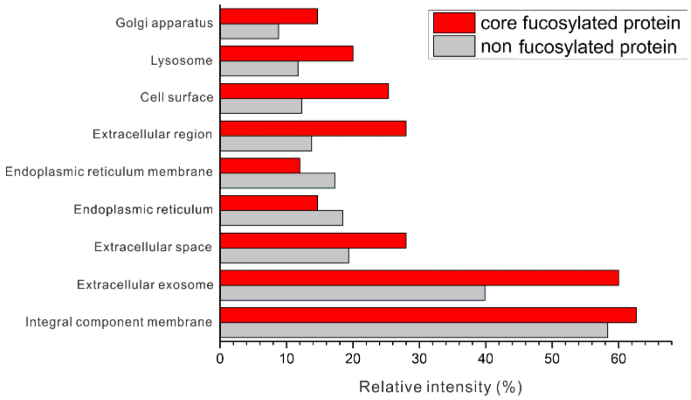


**SFig. 14.** The comparison Cellular compartment distributions of non-fucosylated glycoproteins and core fucosylated glycoproteins.

**SFigure 14**

**STable 1.** The N-glycan detected from MCF 7 and substrates for the endoglycosidases (H, S, F3). YES represents that the glycan could be released by the corresponding endoglycosidase; NO represents that the glycan couldn't be released by the corresponding endoglycosidase

| No. | Chemical composition | m/z | structure | Endo-H | Endo-S | Endo-F3 |
|---|---|---|---|---|---|---|
| 1 | $Hex_3HexNAc_2$ | 565.76 (2+) | 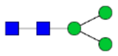 | NO | YES | YES |
| 2 | $Hex_4HexNAc_2$ | 646.68 (2+) | 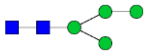 | NO | NO | NO |
| 3 | $Hex_3HexNAc_3$ | 667.30 (2+) | 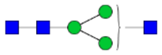 | NO | YES | NO |
| 4 | $Hex_4HexNAc_2$ $DeoxyHex_1$ | 719.8 (2+) | 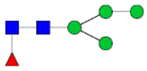 | NO | NO | NO |
| 5 | $Hex_5HexNAc_2$ | 727.81 (2+) | 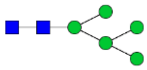 | YES | NO | NO |
| 6 | $Hex_3HexNAc_3$ $DeoxyHex_1$ | 740.33 (2+) | 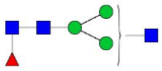 | NO | YES | YES |
| 7 | $Hex_4HexNAc_3$ | 748.32 (2+) | 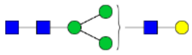 | NO | YES | NO |
| 8 | $Hex_3HexNAc_4$ | 768.84 (2+) | 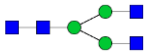 | NO | YES | NO |
| 9 | $Hex_6HexNAc_2$ | 808.84 (2+) | 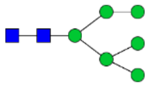 | YES | NO | NO |
| 10 | $Hex_4HexNAc_3$ $DeoxyHex_1$ | 821.35 (2+) | 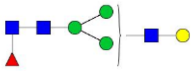 | NO | YES | YES |
| 11 | $Hex_3HexNAc_4$ $DeoxyHex_1$ | 841.87 | 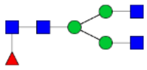 | NO | YES | YES |
| 12 | $Hex_4HexNAc_4$ | 849.86 | 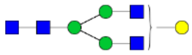 | NO | YES | NO |
| 13 | $Hex_7HexNAc_2$ | 889.86 (2+) | 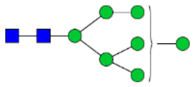 | YES | NO | NO |
| 14 | $Hex_4HexNAc_3Neu5Ac_1$ | 893.87 (2+) | 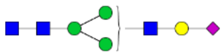 | NO | YES | NO |

| | | | | | | |
|---|---|---|---|---|---|---|
| 15 | Hex$_5$HexNAc$_3$ DeoxyHex$_1$ | 902.38 (2+) | 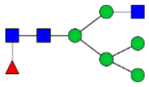 | YES | NO | NO |
| 16 | Hex$_6$HexNAc$_3$ | 910.38 (2+) | 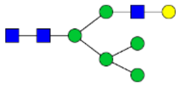 | YES | NO | NO |
| 17 | Hex$_4$HexNAc$_4$ DeoxyHex$_1$ | 922.89 (2+) | 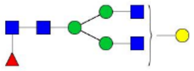 | NO | YES | YES |
| 18 | Hex$_5$HexNAc$_4$ | 930.89 (2+) | 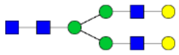 | NO | YES | NO |
| 19 | Hex$_3$HexNAc$_5$ DeoxyHex$_1$ | 943.41 (2+) | 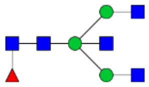 | NO | YES | NO |
| 20 | Hex$_4$HexNAc$_5$ | 951.40 (2+) | 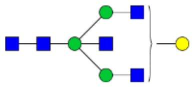 | NO | YES | NO |
| 21 | Hex$_4$HexNAc$_4$Neu5Ac$_1$ | 955.41 (2+) | 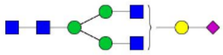 | NO | YES | NO |
| 22 | Hex$_4$HexNAc$_3$Neu5A$_1$ DeoxyHex$_1$ | 966.90 （2+） | 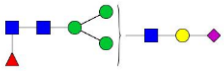 | NO | YES | YES |
| 23 | Hex$_8$HexNAc$_2$ | 970.89 (2+) | 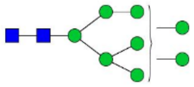 | YES | NO | NO |
| 24 | Hex$_5$HexNAc$_3$Neu5Ac$_1$ | 974.90 (2+) | 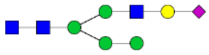 | NO | YES | NO |
| 25 | Hex$_5$HexNAc$_4$ DeoxyHex$_1$ | 1003.91 (2+) | 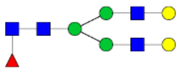 | NO | YES | YES |
| 26 | Hex$_4$HexNAc$_5$ DeoxyHex$_1$ | 1024.43 (2+) | 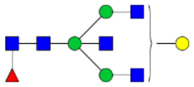 | NO | YES | NO |
| 27 | Hex$_5$HexNAc$_5$ | 1032.42 (2+) | 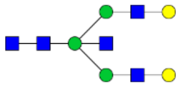 | NO | YES | NO |
| 28 | Hex$_9$HexNAc$_2$ | 1051.92 (2+) | 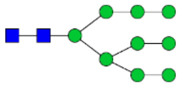 | YES | NO | NO |
| 29 | Hex$_6$HexNAc$_3$Neu5Ac$_1$ | 1055.93 (2+) | 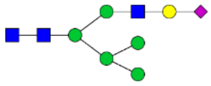 | YES | NO | NO |
| 30 | Hex$_5$HexNAc$_4$Neu5Ac$_1$ | 1076.43 | 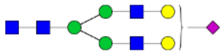 | NO | YES | NO |

| | | | | | | |
|---|---|---|---|---|---|---|
| | | (2+) | | | | |
| 31 | $Hex_5HexNAc_4$ $DeoxyHex_2$ | 1076.94 (2+) | 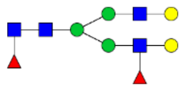 | NO | YES | YES |
| 32 | $Hex_4HexNAc_5Neu5Ac_1$ | 1096.95 (2+) | 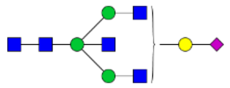 | NO | YES | NO |
| 33 | $Hex_6HexNAc_5$ | 1113.45 (2+) | 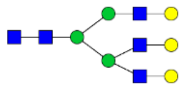 | NO | NO | YES |
| 34 | $Hex_{10}HexNAc_2$ | 1132.94 (2+) | 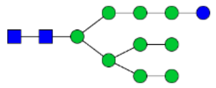 | YES | NO | NO |
| 35 | $Hex_5HexNAc_5Neu5Ac_1$ | 1177.97 (2+) | 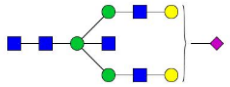 | NO | YES | NO |
| 36 | $Hex_5HexNAc_5$ $DeoxyHex_1$ | 737.30 (3+) | 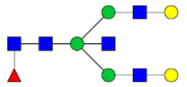 | NO | YES | NO |
| 37 | $Hex_5HexNAc_4Neu5Ac_1$ $DeoxyHex_1$ | 766.65 (3+) | 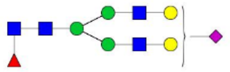 | NO | YES | YES |
| 38 | $Hex_6HexNAc_5$ $DeoxyHex_1$ | 791.33 (3+) | 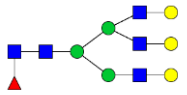 | NO | NO | YES |
| 39 | $Hex_5HexNAc_4Neu5Ac_2$ | 814.99 (3+) | 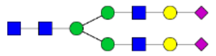 | NO | YES | NO |
| 40 | $Hex_5HexNAc_4Neu5Ac_1$ $DeoxyHex_2$ | 815.33 (3+) | 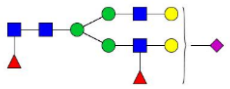 | NO | YES | YES |
| 41 | $Hex_5HexNAc_5Neu5Ac_1$ $DeoxyHex_1$ | 834.98 (3+) | 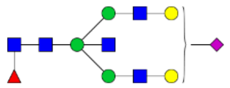 | NO | YES | NO |
| 42 | $Hex_6HexNAc_5Neu5Ac_1$ | 839.67 (3+) | 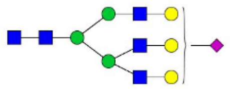 | NO | NO | YES |
| 43 | $Hex_6HexNAc_5$ $DeoxyHex_2$ | 840.01 (3+) | 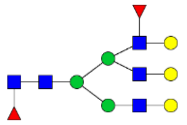 | NO | NO | YES |
| 44 | $Hex_5HexNAc_4Neu5Ac_2$ $DeoxyHex_1$ | 863.67 (3+) | 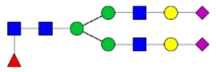 | NO | YES | YES |

| | | | | | | |
|---|---|---|---|---|---|---|
| 45 | $Hex_5HexNAc_5Neu5Ac_2$ | 882.68 (3+) | 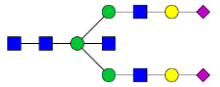 | NO | YES | NO |
| 46 | $Hex_6HexNAc_5Neu5Ac_1$ $DeoxyHex_1$ | 888.35 (3+) | 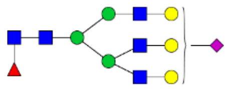 | NO | NO | YES |
| 47 | $Hex_7HexNAc_6$ $DeoxyHex_1$ | 913.03 (3+) | 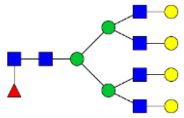 | NO | NO | NO |
| 48 | $Hex_6HexNAc_5Neu5Ac_2$ | 936.70 (3+) | 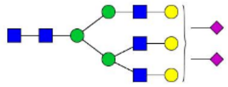 | NO | NO | YES |
| 49 | $Hex_6HexNAc_5Neu5Ac_1$ $Neu5Gc_1$ | 942.03 (3+) | 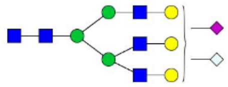 | NO | NO | YES |
| 50 | $Hex_7HexNAc_6Neu5Ac_1$ | 961.38 (3+) | 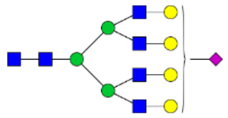 | NO | NO | NO |
| 51 | $Hex_6HexNAc_5Neu5Ac_3$ | 1033.7 3 (3+) | 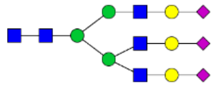 | NO | NO | YES |
| 52 | $Hex_6HexNAc_5Neu5Ac_2$ $DeoxyHex_2$ | 1034.0 7 (3+) | 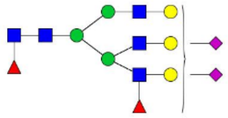 | NO | NO | YES |
| 53 | $Hex_7HexNAc_6Neu5Ac_2$ | 1058.4 1 (3+) | 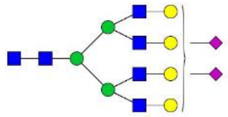 | NO | NO | NO |
| 54 | $Hex_6HexNAc_5Neu5Ac_3$ $DeoxyHex_1$ | 1082.4 2 (3+) | 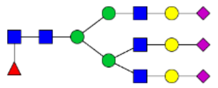 | NO | NO | YES |
| 55 | $Hex_7HexNAc_6Neu5Ac_2$ $DeoxyHex_1$ | 1660.1 4 (2+) | 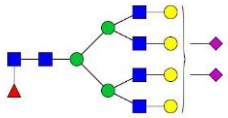 | NO | NO | NO |

**STable 1**

**STable 2** the N-glycan detected from IgG1 Fc fragments, and substrates for the endoglycosidase (H, S, F3). YES represents that the glycan could be released by the corresponding endoglycosidase; NO represents that the glycan couldn't be released by the corresponding endoglycosidase.

| No. | Chemical composition | m/z | structure | Relative intensity (%) | Endo-H | Endo-S | Endo-F3 |
|---|---|---|---|---|---|---|---|
| 1 | $Hex_3HexNAc_3DeoxyHex_1$ | 1831.6 | 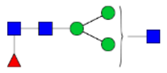 | 2.043 | NO | YES | YES |
| 2 | $Hex_3HexNAc_4$ | 1888.6 | 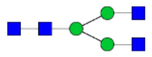 | 1.74 | NO | YES | NO |
| 3 | $Hex_4HexNAc_3DeoxyHex_1$ | 1993.7 | 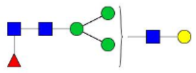 | 1.075 | NO | YES | YES |
| 4 | $Hex_3HexNAc_4DeoxyHex_1$ | 2034.7 | 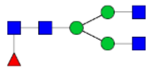 | 20.02 | NO | YES | YES |
| 5 | $Hex_4HexNAc_4$ | 2050.7 | 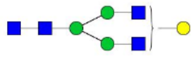 | 4.009 | NO | YES | NO |
| 6 | $Hex_3HexNAc_5$ | 2091.7 | 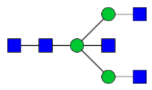 | 0.416 | NO | YES | NO |
| 7 | $Hex_4HexNAc_4DeoxyHex_1$ | 2196.8 | 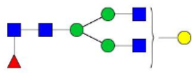 | 29.12 | NO | YES | YES |
| 8 | $Hex_5HexNAc_4$ | 2212.7 | 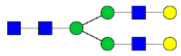 | 2.216 | NO | YES | NO |
| 9 | $Hex_3HexNAc_5DeoxyHex_1$ | 2237.8 | 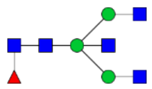 | 4.454 | NO | YES | NO |
| 10 | $Hex_4HexNAc_5$ | 2253.8 | 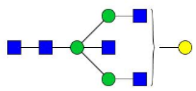 | 0.523 | NO | YES | NO |
| 11 | $Hex_5HexNAc_4Deoxy$ | 2358.8 | 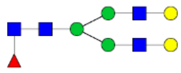 | 14.62 | NO | YES | YES |

| | | | | | | | |
|---|---|---|---|---|---|---|---|
| | $Hex_1$ | | | | | | |
| 12 | $Hex_4HexNAc_5Deoxy$ $Hex_1$ | 2399.8 | 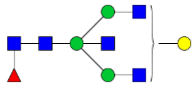 | 5.86 | NO | YES | NO |
| 13 | $Hex_3HexNAc_4Neu5A$ $c_1DeoxyHex_1$ | 2500.9 | 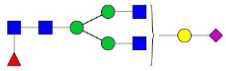 | 2.335 | NO | YES | YES |
| 14 | $Hex_5HexNAc_4Neu5A$ $c_1$ | 2516.9 | 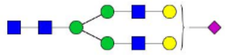 | 0.59 | NO | YES | NO |
| 15 | $Hex_5HexNAc_5Deoxy$ $Hex_1$ | 2561.9 | 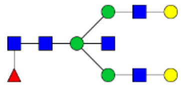 | 1.232 | NO | YES | NO |
| 16 | $Hex_5HexNAc_4Neu5A$ $c_1DeoxyHex_1$ | 2662.9 | 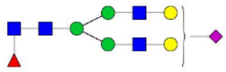 | 9.12 | NO | YES | YES |
| 17 | $Hex_4HexNAc_5Neu5A$ $c_1DeoxyHex_1$ | 2704.0 | 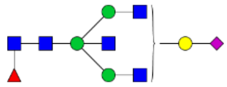 | 0.283 | NO | YES | NO |
| 18 | $Hex_5HexNAc_5Neu5A$ $c_1DeoxyHex_1$ | 2866.0 | 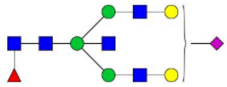 | 0.247 | NO | YES | NO |
| 19 | $Hex_5HexNAc_4Neu5A$ $c_2DeoxyHex_1$ | 2967.1 | 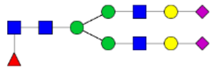 | 0.106 | NO | YES | YES |

**STable 2**
